# Survival Analysis and Prognostic Factors of Time to First Domestic Violence after Marriage among Married Women in Africa

**DOI:** 10.1101/524637

**Authors:** A. F. Fagbamigbe, A. O. Akintayo, O. Oshodi, F. T. Makinde, M. Babalola, E. A. Damilola, O. C. Enabor

## Abstract

**Background:** Domestic violence remains a public health challenge worldwide. Studies in the sub-Saharan Africa have shown high level of violence against women, especially from intimate partners. What is not known is how soon domestic violence happens after marriage among ever-married women. This study assessed the timing of first domestic violence among ever-married women and identified its determinants in Nigeria, Kenya, and Mozambique.

**Data and Method:** Data of 56440 ever-married women of reproductive age pooled from DHS conducted in Nigeria (2013), Kenya (2014) and Mozambique (2011) was used. The timing of first domestic violence was estimated as the period between marriage and the first experience of domestic violence. Ever-married women without violence experience were censored on the date of the survey. Survival analysis techniques were used to assess the timing and the factors influencing the timing p=0.05.

**Result:** The prevalence of domestic violence in Nigeria, Kenya and Mozambique was 12.1%, 7.5%, and 15.4% respectively. The median time to first domestic violence across the three countries was 3 years. We found a higher prevalence of violence among formerly married women than currently married women. Women who married before age 20 years have a significantly higher risk of experiencing violence (Nigeria: aHR= 2.36 (95% CI (1.97- 2.83), Kenya: aHR= 1.89 (95% CI (1.31- 2.73) and Mozambique: aHR=1.60 (95% CI (1.24 −2.12)) than older women. Women in poorer wealth quintile were at higher risk of violence across the three countries. Other factors associated with the timing of first domestic violence are respondents’, location of residence and educational attainment.

**Conclusion:** Domestic violence has remained high in Mozambique, Nigeria and Kenya. There was a significant relationship between timing of first domestic violence and whether or not a woman remains in a marriage. Education and delayed marriage could help reduce violence in marriage.

## BACKGROUND

Violence is a fundamental human right issue and has constituted global social, clinical health as well as public health challenges(1–4). Literature is replete that violence against women, especially women and girls has remained high despite several interventions to halt it(5,6). Depending on the presence or absence of a relationship between a victim and a perpetrator of a violence, violence can be grouped as self-directed, interpersonal and collective violence. Domestic violence is one of the interpersonal violence and has unacceptably thrived across various regions of the world. Domestic violence, also referred to as spousal or intimate partner violence, is a subtle type of violence occurring between partners in union (legal or not) and within the family context. Domestic violence happens both in cohabitation and marriage. Nevertheless, occurrence may not be restricted to the confines of the home(3). Domestic violence is not peculiar to only heterosexuals, it occurs in both same and different-sex marriages and is perpetrated by both men and women. However, research has overtime revealed that domestic violence are mostly perpetrated by men against the women.

Despite the development of gender policy in Nigeria and other African countries to promote the health and protect the rights of women, domestic violence remains a menace to the health and wellbeing of African women(7). This is fostered among other factors by unhealthy sociocultural norms that presupposes African women as inferior to men(8). Sadly, significant proportion of women in Africa hold perceptions which justify physical assault of a woman by her husband or partner, thereby encouraging under-reporting of spousal violence. The level of domestic violence in Kenya has been established by various studies with physical violence and sexual prevalence placed at 42% and 14% respectively. The figures lie in the middle range of the WHO multi-country estimate 14-61% and 6-59% respectively(9,10).

The prevalence of domestic violence is diverse across different settings. While ample evidence exists that supports the occurrence of domestic violence across the world, disparity exists in its prevalence among low and middle income countries (LMIC) as well as high income countries. For instance, 15% to 71% of ever-married women have been reported to have been abused either physically or sexually by an intimate partner throughout at some points in their lives(11). In a multi-regional study carried out by WHO, the prevalence of lifetime sexual and/or physical spousal violence among ever-married women was 30%. The highest prevalence was found in the low and middle income countries among ever-married women in the WHO South-East Asia, Eastern Mediterranean and African regions (37.7%, 37.0% and 36.6%) compared to 23.2% in high income countries(10).

Multiple studies have reported increased occurrence of violence against women in the sub-Saharan Africa region, particularly in Nigeria which is the largest country in Africa(12–14). A recent study found a prevalence of spousal violence (physical, sexual or emotional) of about 30.5% in Nigeria 45.3% in Kenya and 45.5% in Mozambique(4). In a meta-analysis of 141 studies conducted in 81 countries the frequency of lifetime exposure to domestic violence among women was found to be relatively high with 65.6% in central sub-Saharan Africa, western sub-Saharan Africa (41.8%) and South Asia (41.7%)(15).

The consequence of domestic violence on women is quite colossal. Women’s sexual and reproductive health suffers incessant denting(16). Besides maternal depletion disorder in conjunction with its associated problems, research has shown that inadvertent pregnancy termination can also be linked to domestic violence among married persons(8,17). Furthermore, research in the past decades has provided sufficient evidence that suicide, homicide, physical injuries, increased susceptibility to sexually transmitted infections, unwanted pregnancies, forced abortion, gynecological complaints, low body mass index infant and child mortality are possible adverse effects of spousal violence on women’s reproductive health(5,7,18–21). Physical and sexual violence by partner has been linked with increased transmission of HIV among Kenyan women via tears and laceration of the vaginal canal(22,23). In Nigeria, domestic violence has been linked to an increase in the incidence of miscarriages, induced abortions and stillbirths(24).

Literature have identified risk factors of domestic violence to include age at marriage or cohabitation, religion, education, place of residence and household wealth quintile and employment(4,10,12,19). A recent multi-regional study by WHO showed that the prevalence of domestic violence among ever-married women was about 29.4% in women aged 15-19 years, suggesting that IPV starts early in life, but higher (over 36%) among older women(10). Another multi-country study in sub-Saharan Africa linked higher prevalence of violence to higher household wealth quintile in Mozambique while the reverse was the case for Kenya as women from poor households were reported to be more probable suffer domestic violence(4).

While the differential pattern and distribution of domestic violence among married and cohabiting women has been evaluated(4,25–29), the timing of the occurrence of first domestic violence has not been explored in sub-Saharan Africa. Similarly, study outcomes abound on the prevalence of domestic violence and the factors influencing it(4,26–29), but there remains no information on the factors that could affect the risk and the time to first domestic violence among ever-married women in sub-Saharan Africa. The current study is therefore designed to assess how long it takes married women in Nigeria, Kenya and Mozambique Africa to first experience domestic violence and also to identify the risk factors of the timing. Considerably, new insights provided by this paper would be useful for effective intervention against spousal abuse in Nigeria, Kenya and Mozambique and possibly sub-Saharan Africa.

## Materials and Methods

### Study Setting

This study used data from the most recent sets of Demographic and Health Survey (DHS) conducted in Nigeria (2013), Kenya (2014), and Mozambique (2011) conducted by ICF Macro Calverton, Maryland, USA in conjunction with Population Councils of three different African countries. All the DHSs were cross-sectional and nationally representative. The surveys used the same set of questions to obtain information on demographics, sexuality and other reproductive and sexual behavior of men and women of reproductive age. All the DHS used in this study provided population and health indicator estimates at the national and provincial/district/state levels.

### Sampling and Samples Nigeria

The Nigeria DHS 2013 adopted probability multistage sampling procedure to select representative sample of women. The sampling used the enumeration areas (EA) that were prepared from the 2006 population census of Nigeria as primary sampling units. The sample was selected using a stratified three-stage cluster design consisting of 904 clusters—372 in urban areas and 532 in rural areas. At the first stage, local governments were selected, then EAs were selected at the second stage while the households were selected at the third stage. A representative sample of 40 680 households was selected for the survey among which 38948 were successfully interviewed. Detailed sampling details have been documented (30).

### Kenya

The sample for the 2014 Kenya DHS was drawn from the Fifth National Sample Survey, and Evaluation Programme (NASSEP-V). Kenya is divided into 47 counties with each county stratified into urban and rural strata (clusters). Using a two-stage sampling design, samples were drawn in each stratum independent of other strata. In the first stage, 1612 clusters were randomly selected with equal probability selection method from the NASSEP-V frame, with 617 from urban and 995 from rural areas. Twenty-five households were randomly selected from each cluster using the list of eligible households at the second stage. A total of 40,300 households were visited among which 31,079 women were interviewed.

### Mozambique

The sample for the 2011 Mozambique DHS was selected from the 11 provinces in the country. Each province was divided in districts, and the districts further subdivided into administrative posts using the master sample designed by the National Statistical Office and the US Census Bureau using the 1997 household and population census. A representative probability sample of approximately 14,500 households was selected for the 2011 MDHS among which 13745 were successfully interviewed. Samples of districts were drawn at the first stage within each district, samples of administrative polls drawn at the second stage from the selected districts, while the EAs and the households were subsequently drawn at the 3^rd^ and 4^th^ stages.

### Sampling of respondents on Violence related questions

Due to the sensitivity of violence and to maintain confidentiality, only one woman per household was administered the questions on violence while. In the one-third of the households selected for the male survey, one man per household was randomly selected to respond to questions on domestic violence. One woman per household was randomly selected in the remaining two-thirds of households and were administered the questions on violence. The selected respondents were informed that questions could be sensitive and were reassured regarding the confidentiality of their responses.

### Data

To determine whether an ever-married woman had experienced domestic violence prior to the survey dates, the following questions were asked. Were you: ever been pushed, shook or had something thrown by husband/partner, ever been slapped by husband/partner, ever had arm twisted or hair pulled by husband/partner; ever been punched with fist or hit by something harmful by husband/partner, ever been kicked or dragged by husband/partner; ever been strangled or burnt by husband/partner ever been threatened with knife/gun or other weapon by husband/partner; ever been physically forced into unwanted sex by husband/partner; ever been forced into other unwanted, sexual acts by husband/partner; ever been physically forced to perform sexual acts respondent didn’t want to.

Any ever-married who answered in affirmative to at least one of the question is considered to have experienced domestic violence and were then asked how long was the first time they experienced any form of these violence after they got married. The length of time between marriage date and the date of first domestic violence was used as the dependent variable in this study while age, education, employment, residence and wealth indicators were used as independent variable. They have been identified in prior studies as contributing factors to domestic violence among women(4,10,12,19).

### Inclusion and exclusion criteria

The current study focused on experience of domestic violence among ever-married women. We therefore excluded never married women either they were living with sexual partner or not. Only women aged 15-49 years and who were currently married, divorced, separated or widowed were included in the analysis. The effective sampling size for this study was 21,564, 4,237, and 3992 for Nigeria, Kenya and Mozambique respectively totaling 29,793 respondents. Of the 6440 that claimed to have experienced violence before, 343 did not remember their time of first experience after marriage, they were therefore excluded in further analysis except in Table 1.

**Table 1:** Distribution of ever-married women by socio-demographic characteristics and violence experience across the selected countries.

| Characteristics | <u>Distribution of Respondents (%)</u> |  |  |  | <u>Prevalence of Domestic Violence (%)</u> |  |  |  |
| --- | --- | --- | --- | --- | --- | --- | --- | --- |
|  | NIG | KEN | MOZ | All | NIG | KEN | MOZ | All |
| **Age** |  |  |  |  |  |  |  |  |
| 15-19 | 7.7 | 2.8 | 10.9 | 7.7 | 9.3 | 22.4 | 22.6 | 13.4 |
| 20-24 | 15.5 | 16.1 | 19.2 | 16.2 | 14.6 | 36.5 | 32.7 | 21.3 |
| 25-34 | 37.5 | 43.8 | 35.6 | 38.0 | 16.2 | 38.8 | 33.3 | 22.5 |
| 35-49 | 39.3 | 37.3 | 34.3 | 38.1 | 16.1 | 41.4 | 30.3 | 21.6 |
| Marital Status |  |  |  |  |  |  |  |  |
| Currently Married | 93.8 | 82.0 | 76.4 | 89.8 | 14.0 | 36.1 | 25.7 | 18.0 |
| Widowed | 3.4 | 5.8 | 7.0 | 4.2 | 17.5 | 42.7 | 24.0 | 23.6 |
| Separated/Divorced | 2.8 | 12.2 | 16.6 | 6.0 | 46.7 | 61.4 | 44.2 | 49.8 |
| Residence |  |  |  |  |  |  |  |  |
| Urban | 37.2 | 39.5 | 30.7 | 36.3 | 16.8 | 35.8 | 34.3 | 22.2 |
| Rural | 62.8 | 60.5 | 69.3 | 63.7 | 14.5 | 41.0 | 29.5 | 20.8 |
| Education |  |  |  |  |  |  |  |  |
| No education | 47.1 | 9.3 | 36.2 | 40.2 | 9.0 | 33.1 | 28.6 | 12.9 |
| Primary | 19.7 | 56.4 | 50.8 | 30.2 | 24.1 | 44.0 | 32.9 | 31.7 |
| Secondary | 28.8 | 26.0 | 12.0 | 23.3 | 21.4 | 35.2 | 30.4 | 24.3 |
| Tertiary | 7.4 | 8.3 | 1.0 | 6.3 | 12.1 | 22.7 | 26.8 | 14.3 |
| Husband education |  |  |  |  |  |  |  |  |
| No education | 39.1 | 7.5 | 24.9 | 32.5 | 8.46 | 37.1 | 29.4 | 12.1 |
| Primary | 18.4 | 47.3 | 53.9 | 28.3 | 22.1 | 43.5 | 32.0 | 30.1 |
| Secondary | 28.5 | 33.1 | 19.2 | 27.5 | 20.8 | 37.8 | 31.0 | 24.7 |
| Tertiary | 14.0 | 12.2 | 2.0 | 11.7 | 15.1 | 26.3 | 20.8 | 16.7 |
| Employment |  |  |  |  |  |  |  |  |
| Not employed | 28.1 | 23.3 | 48.7 | 31.2 | 10.8 | 25.2 | 28.7 | 17.2 |
| Employed | 71.9 | 76.7 | 51.3 | 68.8 | 17.2 | 43.0 | 33.1 | 23.1 |
| Husband employment |  |  |  |  |  |  |  |  |
| Not employed | 1.2 | 1.6 | 4.3 | 1.8 | 20.7 | 36.1 | 33.1 | 27.8 |
| Employed | 98.8 | 98.4 | 95.7 | 98.2 | 15.3 | 39.1 | 30.9 | 21.2 |
| Wealth Category |  |  |  |  |  |  |  |  |
| Poorest | 22.2 | 17.6 | 20.0 | 21.2 | 9.8 | 39.7 | 31.4 | 16.7 |
| Poorer | 21.2 | 19.4 | 20.3 | 20.8 | 15.1 | 46.8 | 27.7 | 21.2 |
| Middle | 18.5 | 18.9 | 21.2 | 19.1 | 19.4 | 42.5 | 29.8 | 24.5 |
| Richest | 18.7 | 22.2 | 20.3 | 19.4 | 17.1 | 37.0 | 33.3 | 23.1 |
| Richest | 19.4 | 21.9 | 18.2 | 19.5 | 16.7 | 30.8 | 33.1 | 21.4 |
| Age difference |  |  |  |  |  |  |  |  |
| Wife Older | 1.0 | 3.0 | 5.6 | 2.0 | 27.9 | 30.3 | 22.2 | 25.7 |
| Same age | 1.4 | 3.3 | 3.4 | 1.9 | 14.1 | 32.4 | 37.7 | 24.6 |
| Wife younger | 97.6 | 93.7 | 91.0 | 96.1 | 14.3 | 35.8 | 29.6 | 19.2 |
| Husband takes |  |  |  |  |  |  |  |  |
| No | 81.6 | 63.3 | 61.1 | 75.5 | 10.7 | 29.8 | 23.1 | 14.6 |
| Yes | 18.4 | 36.7 | 38.9 | 24.5 | 37.0 | 54.7 | 43.5 | 42.4 |
| Husband Jealous |  |  |  |  |  |  |  |  |
| No | 42.5 | 46.8 | 43.3 | 43.2 | 9.9 | 23.8 | 21.1 | 13.4 |
| Yes | 57.5 | 53.2 | 56.7 | 56.8 | 19.5 | 52.3 | 32.3 | 27.3 |
| Total | 21564 | 4237 | 3992 | 29793 | 15.4 | 39.0 | 31.0 | 21.3 |
| NIG=Nigeria, KEN= Kenya, MOZ=Mozambique. |  |  |  |  |  |  |  |  |

### Data Analysis

Descriptive statistics and survival analysis techniques were used to analyze the data at a 5% significance level. Cox proportional hazard regression model was used to identify the risk factors associated with the timings. All data were weighed so as to ensure adequate representativeness at the national, regional, and county levels because of the non-proportional allocation to the sampling strata and the fixed sample size per cluster. The statistical analysis was carried out using IBM SPSS 24.

### Justification for use of Survival Analysis

Right from the day of marriage, there are possibilities that an ever-married woman is violated by her husband. However, it is not impossible that there are ever-married women without any experience of domestic violence as of the survey date. Non-inclusion of ever-married women who have not experience any domestic violence as of the day of the survey might seriously bias the computation of the timing of first domestic violence after marriage as well as its risk factor. The survival analysis remains the utmost data analysis procedure for lifetime data when some subjects have not experience the event of interest. Therefore, the populations at risk is all married women involved in the study. The duration from marriage to first domestic violence, ‘T’, is assumed to be a discrete random variable that is always positive. The observation continues until time ‘t’, at which the event of interest (first domestic violence) occurs. The study ends for an individual at time ‘T=t’ if she experienced any form of domestic violence after marriage. Survival analysis requires the censoring index and the survival time as basic variables. The survival time is assumed to begin at the time a woman had her first marriage until the time she experienced first domestic violence after marriage. The survival time in our study is the length of time (years) between the date of marriage and the date of first domestic violence by the husband, for those who had experienced violence. For the women who did not experience any domestic violence as of survey date, the survival time is censored and the age of their marriage in years is used as their survival times.

The Kaplan Meier techniques were used to estimate survival and hazard functions. The survivor function is the probability that a woman survives longer than some specified time t without experience any form of domestic violence, while the hazard function is the instantaneous potential per unit of time for domestic violence to occur, given that the woman had not experience any domestic violence before time t.

We used the Cox proportional-hazards to model risk factors of the timings and estimate the strength of the relationship between each of the selected independent variables and censored timing of first domestic violence. The outcomes of the model are the hazard ratios. A hazard ratio (HR) <1 means higher risk, <1 means lower risk while =1 suggest insignificantly different risk.

We tested whether the proportional-hazards assumption was violated using the significance of the HRs and Wald χ2 statistics in our stratified Cox analysis. We moved the significant variables in the bivariate Cox regression models into the multiple Cox regression to assess their association with the outcome variable while controlling for confounders.

## RESULTS

Table 1 shows the distribution of the ever-married women by their socio-demographic characteristics and the prevalence of ever-experience of domestic violence across the three countries. In all 89.8% were currently married, almost a two-third (63.7%) were from rural areas, 30.2% had had primary education, 31.2% were unemployed while 7.7%, 16.2%, 38.0%, and 38.1% were aged 15-19, 20-24, 25-34 and 35-49 years respectively.

In Nigeria, 93.8% of the ever-married women were currently married, about two-thirds (62.8%) were from rural areas, almost half of the respondents (47.10%) had had no formal education, while 98.8% were employed. Of the 4237 ever-married women in Kenya, 82.0% were currently married, about two-thirds (60.5%) were from rural areas, more than half (56.4%) had had primary education, 23.3% were not working, 2.8%, 16.1%, 43.8%, and 37.3% were aged 15-19, 20-24, 25-34 and 35-49 years respectively. Over 76% ever-married women in Mozambique were currently married, about two-thirds (69.3%) were from rural areas, 50.8% had had primary education, 48.7% were not working while 10.9%, 19.2%, 35.6%, and 34.3% were aged 15-19, 20-24, 25-34 and 35-49 years respectively.

Of the 29,793 ever-married women across the three countries, 6440 (21.3%) reported ever experienced domestic violence. The prevalence of ever-experience of domestic violence was 15.4%, 39.0% and 31.0% in Nigeria, Kenya and Mozambique respectively. In Nigeria, the highest prevalence of domestic violence was among respondents aged 25-34 years (16.2%), 35-49 years (16.1%), 46.7% among separated/divorced women, 24.1% among respondents with primary education, 17.2% among employed women, 27.9% among women who are older than their husbands, 37.0% among women whose husband takes alcohol and 19.5% among women whose husband get jealous when they talk to other men. Also, the highest report of domestic violence in Kenya was found among respondents aged 25-34 years (38.8%), 35-49 years (41.4%), 61.4% among separated/divorced women, 44.0% among respondents with primary education, 43.0% among employed women, 35.8% among women who are of younger than their husbands, 54.7% among women whose husband takes alcohol and 52.3% among women whose husband get jealous when they talk to other men. In Mozambique, the highest prevalence of domestic violence was among respondents aged 25-34 years (33.3%), 20-24 years (32.7%), 44.2% among separated/divorced women, 32.9% among respondents with primary education, 33.1% among employed women, 37.7% among women who are of same age as their husbands, 43.5% among women whose husband takes alcohol and 32.3% among women whose husband get jealous when they talk to other men.

In Table 2, we present the distribution of timing of first domestic violence among ever-married women who have already experienced the act. The median time to first domestic violence was 3 years in Nigeria and Kenya and 2 years in Mozambique. The timing varied across the women characteristics. For instance, 29.6% of the currently married women in Nigeria experienced first domestic violence within one year of marriage compared with 33.5% among the separated/divorced and 12.5% among the widows. In Mozambique, 48.0% of the currently married women experienced first domestic violence within one year of marriage compared with 49.3% among the separated/divorced and 54.8% among the widows within the same period.

**Table 2:** Distribution of time to first domestic violence after marriage among ever-married women in the three countries.

| Characteristics | Nigeria |  |  |  | Kenya |  |  |  | Mozambique |  |  |  |
| --- | --- | --- | --- | --- | --- | --- | --- | --- | --- | --- | --- | --- |
|  | Median years to FDV | Years to FDV (%) |  |  | Median years to FDV | Years to FDV (%) |  |  | Median years to FDV | Years to FDV (%) |  |  |
|  |  | <1 | 2-3 | >3 |  | <1 | 2-3 | >3 |  | <1 | 2-3 | >3 |
| **Age** |  |  |  |  |  |  |  |  |  |  |  |  |
| 15-19 | 1 | 75.2 | 18.9 | 5.9 | 0 | 93.7 | 6.4 | 0.0 | 1 | 75.4 | 18.9 | 5.7 |
| 20-24 | 1 | 56.0 | 29.8 | 14.2 | 2 | 55.7 | 31.0 | 13.3 | 1 | 61.6 | 27.3 | 11.0 |
| 25-34 | 3 | 30.0 | 36.2 | 33.8 | 3 | 32.6 | 34.6 | 32.8 | 2 | 44.2 | 36.0 | 19.8 |
| 35-49 | 4 | 16.7 | 27.1 | 56.2 | 4 | 17.2 | 27.2 | 55.7 | 3 | 36.8 | 28.8 | 34.4 |
| **Marital Status** |  |  |  |  |  |  |  |  |  |  |  |  |
| Currently Married | 3 | 29.6 | 30.2 | 40.2 | 3 | 29.2 | 30.1 | 40.7 | 2 | 48.0 | 32.0 | 20.0 |
| Widowed | 3 | 12.5 | 39.4 | 48.1 | 3 | 27.6 | 33.6 | 38.7 | 2 | 54.8 | 25.6 | 19.6 |
| Separated/Divorced | 2 | 33.5 | 32.0 | 34.6 | 2 | 37.5 | 32.5 | 30.0 | 2 | 49.3 | 32.3 | 18.4 |
| **Residence** |  |  |  |  |  |  |  |  |  |  |  |  |
| Urban | 3 | 26.7 | 34.3 | 39.0 | 2 | 36.8 | 31.0 | 32.2 | 2 | 51.6 | 27.2 | 21.2 |
| Rural | 3 | 33.0 | 28.3 | 38.7 | 3 | 27.6 | 30.5 | 42.0 | 2 | 45.8 | 32.1 | 22.1 |
| **Education** | 3 |  |  |  |  |  |  |  | 2 |  |  |  |
| No education | 3 | 27.6 | 26.6 | 45.9 | 3 | 31.9 | 29.0 | 39.0 | 2 | 40.3 | 34.5 | 25.2 |
| Primary | 3 | 27.7 | 29.9 | 42.4 | 3 | 29.0 | 31.6 | 39.4 | 2 | 51.6 | 28.0 | 20.4 |
| Secondary | 3 | 34.2 | 35.1 | 30.7 | 3 | 32.4 | 30.2 | 37.5 | 2 | 51.0 | 30.4 | 18.6 |
| Tertiary | 3 | 35.3 | 28.5 | 36.2 | 2 | 46.9 | 24.1 | 29.0 | 2 | 58.9 | 26.9 | 14.2 |
| **Husband education** |  |  |  |  |  |  |  |  |  |  |  |  |
| No education | 3 | 25.7 | 26.3 | 48.0 | 2 | 29.5 | 29.1 | 41.4 | 2 | 44.5 | 29.5 | 25.9 |
| Primary | 3 | 29.4 | 29.9 | 40.7 | 3 | 28.5 | 31.9 | 39.5 | 2 | 46.0 | 31.0 | 23.0 |
| Secondary | 3 | 33.0 | 34.4 | 32.5 | 3 | 34.3 | 29.6 | 36.1 | 2 | 52.0 | 30.8 | 17.2 |
| Tertiary | 3 | 32.7 | 28.2 | 39.2 | 3 | 35.8 | 28.2 | 36.0 | 2 | 60.1 | 27.4 | 12.5 |
| **Employment** |  |  |  |  |  |  |  |  |  |  |  |  |
| Not employed | 2 | 39.9 | 27.8 | 32.2 | 2 | 31.6 | 36.3 | 32.1 | 2 | 52.9 | 27.1 | 19.9 |
| Employed | 3 | 28.1 | 31.4 | 40.5 | 3 | 30.6 | 29.9 | 39.6 | 2 | 43.6 | 33.2 | 23.3 |
| **Husband** | 3 |  |  |  |  |  |  |  |  |  |  |  |
| Not employed | 3 | 33.2 | 18.8 | 48.0 | 4 | 39.4 | 14.1 | 46.6 | 2 | 48.9 | 35.4 | 15.6 |
| Employed | 3 | 30.4 | 30.9 | 38.7 | 3 | 30.7 | 31.0 | 38.3 | 2 | 47.7 | 30.2 | 22.1 |
| Wealth Category |  |  |  |  |  |  |  |  |  |  |  |  |
| Poorest | 3 | 24.5 | 30.0 | 45.5 | 3 | 27.6 | 30.1 | 42.3 | 2 | 43.3 | 37.6 | 19.1 |
| Poorer | 3 | 32.4 | 28.9 | 38.8 | 3 | 28.8 | 32.8 | 38.4 | 2 | 46.7 | 30.1 | 23.2 |
| Middle | 3 | 33.9 | 26.5 | 39.7 | 3 | 27.1 | 31.6 | 41.3 | 2 | 51.0 | 30.1 | 18.9 |
| Richest | 3 | 30.6 | 34.6 | 34.8 | 2 | 36.1 | 28.6 | 35.3 | 2 | 47.1 | 27.4 | 25.5 |
| Richest | 3 | 28.6 | 34.0 | 37.4 | 2 | 35.2 | 29.9 | 34.9 | 2 | 51.0 | 27.0 | 22.0 |
| Age difference |  |  |  |  |  |  |  |  |  |  |  |  |
| Wife Older | 3 | 29.9 | 29.8 | 40.3 | 4 | 37.4 | 23.3 | 39.3 | 2 | 55.2 | 27.6 | 17.2 |
| Same age | 3 | 37.5 | 18.0 | 44.4 | 3 | 31.1 | 21.7 | 47.2 | 2 | 69.0 | 23.4 | 7.6 |
| Wife younger | 3 | 30.9 | 30.4 | 38.7 | 3 | 29.0 | 30.6 | 40.4 | 2 | 45.7 | 30.8 | 23.6 |
| Husband takes |  |  |  |  |  |  |  |  |  |  |  |  |
| No | 3 | 31.4 | 29.7 | 38.9 | 3 | 31.4 | 28.5 | 40.1 | 2 | 48.6 | 33.3 | 18.1 |
| Yes | 3 | 29.1 | 32.1 | 38.8 | 3 | 30.5 | 32.7 | 36.9 | 2 | 47.1 | 28.1 | 24.9 |
| Husband Jealous |  |  |  |  |  |  |  |  |  |  |  |  |
| No | 3 | 27.9 | 31.9 | 40.2 | 3 | 29.7 | 25.9 | 44.5 | 2 | 45.9 | 29.5 | 24.6 |
| Yes | 3 | 31.2 | 30.4 | 38.4 | 3 | 31.4 | 32.7 | 35.9 | 2 | 48.5 | 30.9 | 20.6 |
| Total | 3 | 30.4 | 30.7 | 38.8 | 3 | 30.9 | 30.7 | 38.4 | 2 | 47.8 | 30.5 | 21.7 |
| FDV First Forced Sex *Age in years Analysis restricted to 7002 ever married women who had experienced |  |  |  |  |  |  |  |  |  |  |  |  |

The timing also appeared increased with women educational attainment, while the proportion of women who experienced first domestic violence within one year of marriage among those that had no education, primary, secondary and higher was 27.6%, 27.7%, 34.2%, and 35.3% in Nigeria, the figures were 31.9%, 29.0%, 32.4% and 46.9% in Kenya and 40.3%, 51.6%, 51.0% and 58.9% in Mozambique. On the average, 30.4% of ever-women in Nigeria, 30.9% in Kenya and 47.8% in Mozambique had first domestic violence within one year of marriage.

The Figures 1, 2, and 3 show the distribution of probabilities of falling victim to domestic violence as a woman remains in her marriage in the three countries. According to the Log-rank test of equality of survival curves, the timing of first domestic violence differed significantly across the women age, marital status, place of residence, educational attainment, husband educational attainment, employment status, husbands’ employment status, whether husband takes alcohol, gets jealous when the woman talks to another man, and the wealth quintile their households in the three counties except between place of residence in Mozambique (Figures 1, 2, and 3).

**Figure 1:**
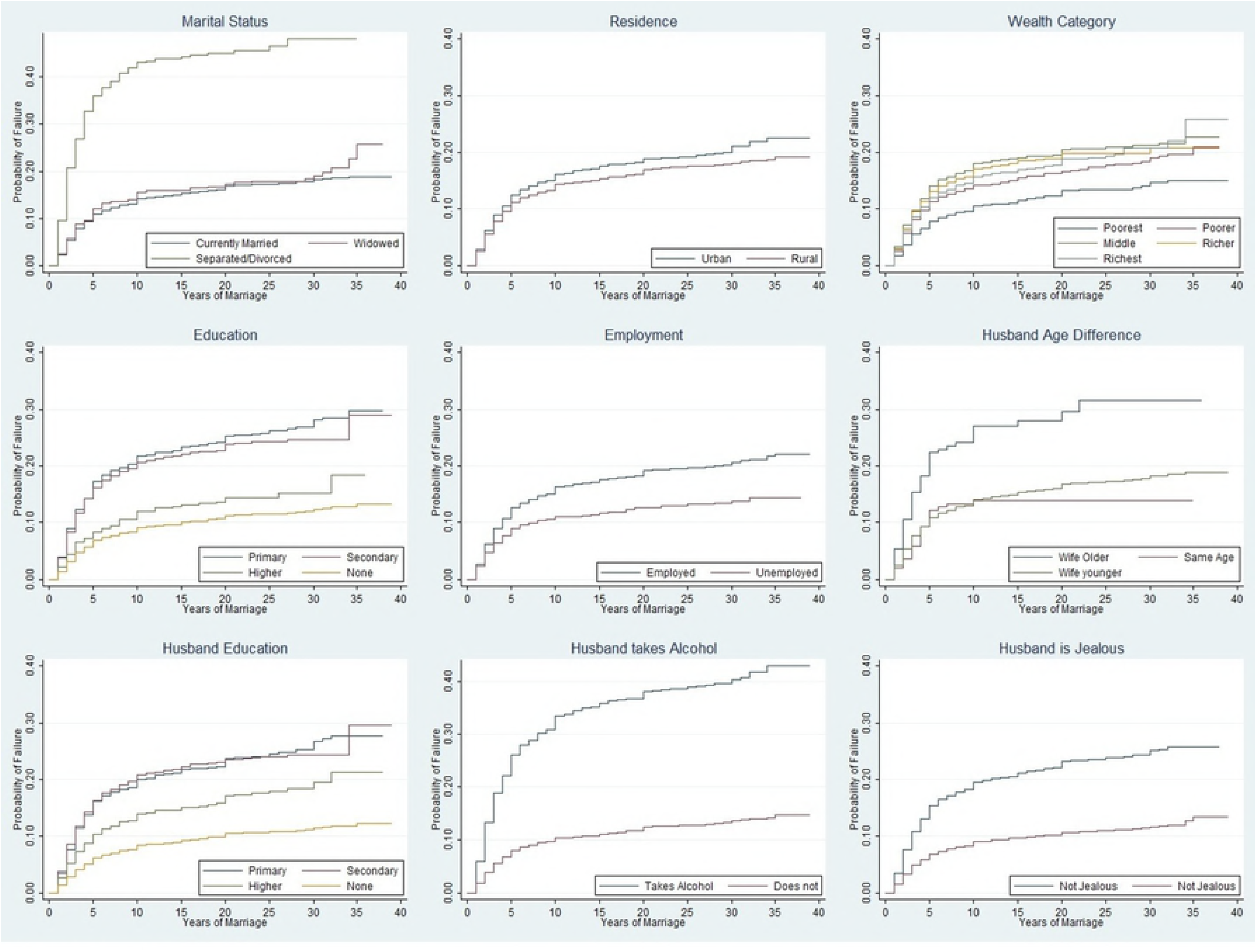
Kaplan Meier Probability Plots of Timing of first domestic violence among ever-married women in Nigeria by selected socio-demographic characteristics.

**Figure 2:**
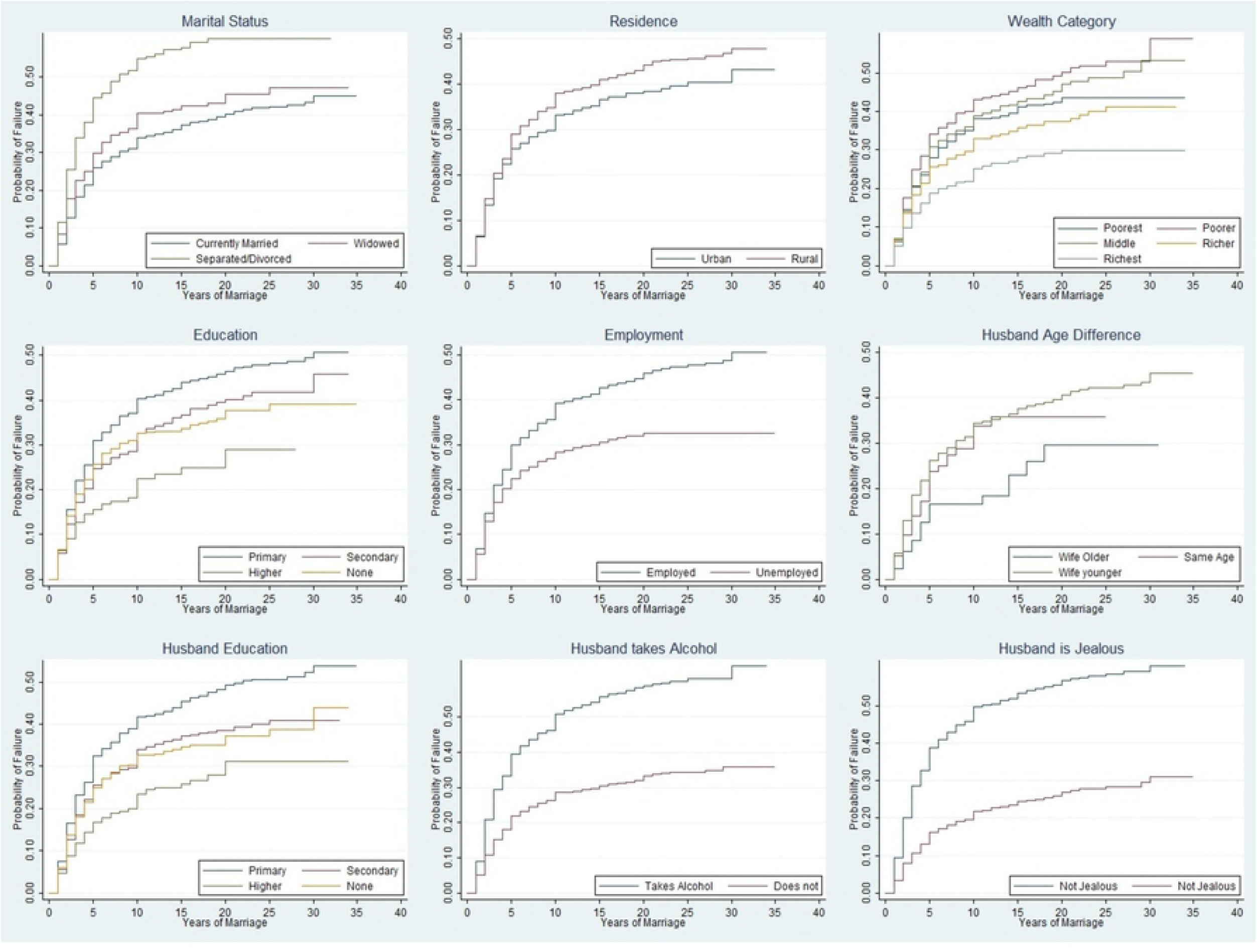
Kaplan Meier Probability Plots of Timing of first domestic violence among ever-married women in Kenya by selected socio-demographic characteristics.

**Figure 3:**
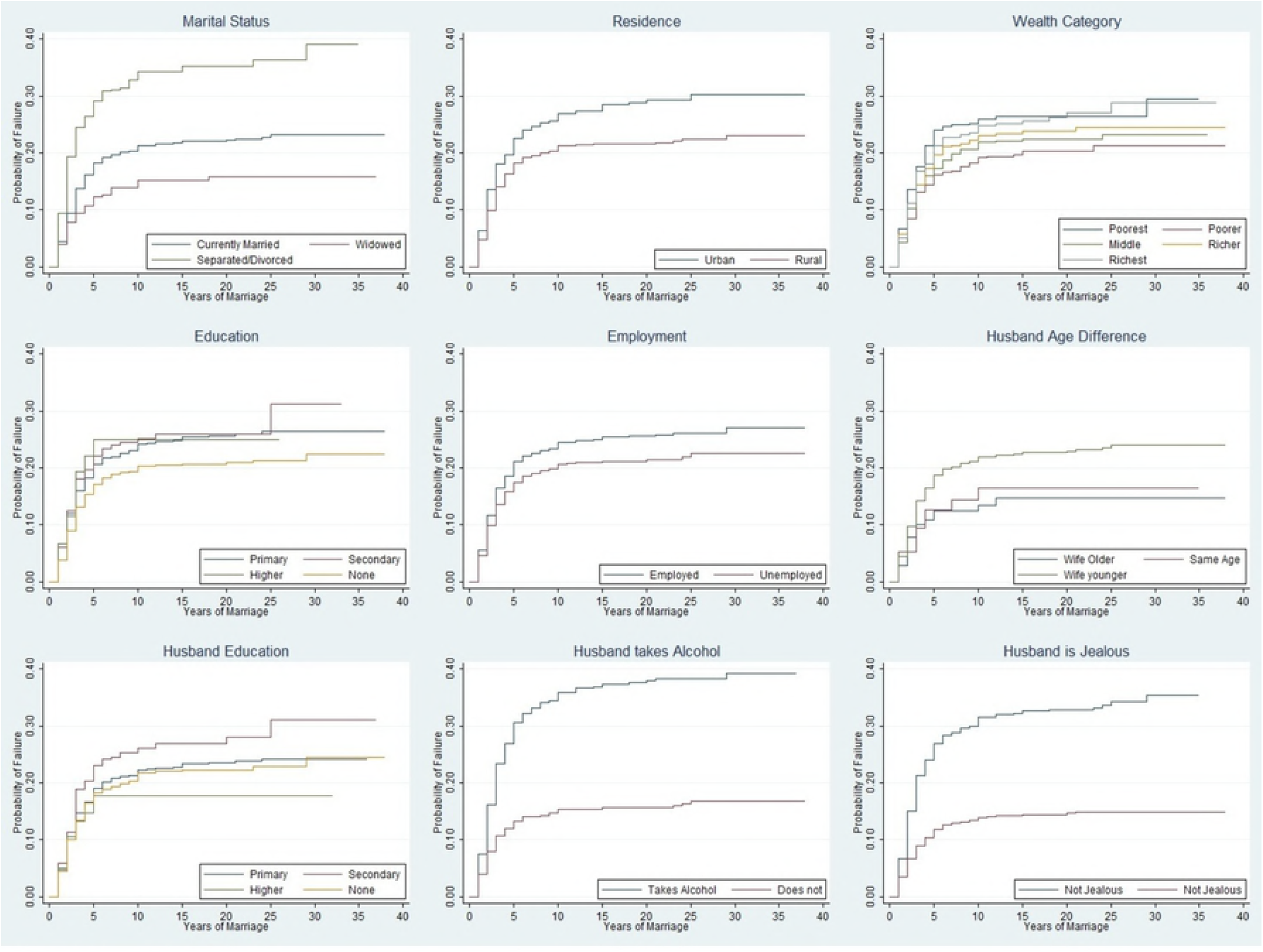
Kaplan Meier Probability Plots of Timing of first domestic violence among ever-married women in Mozambique by selected socio-demographic characteristics.

Table 3 shows the unadjusted (crude) determinants of the timing of first domestic violence after marriage among ever-married women in the three countries. The risk of experiencing domestic violence among ever-married women was significantly higher among separated/divorced and widowed women than currently married women across the three countries (HR= 3.467; 95% CI=3.022-3.979; HR=1.847: 95% CI=1.582-2.154 and HR=1.747: 95% CI=1.469-2.077 for Nigeria, Kenya and Mozambique respectively). The risk of domestic violence was higher among younger ever-married women compared with those aged 35-49 years in the three countries. For instance, the risk among women aged 20-24 years in Nigeria, Kenya and Mozambique were about 50% times higher than the risk among women aged 45-49 years.

**Table 3:** The unadjusted determinants of domestic violence after marriage.

| Characteristics | Unadjusted determinants (HR &95% CI) |  |  |
| --- | --- | --- | --- |
|  | Nigeria | Kenya | Mozambique |
| **Age** |  |  |  |
| 15-19 | 1.196(0.972-1.470) | 1.227(0.744-2.024) | 1.392(1.066-1.817) |
| 20-24 | 1.311(1.167-1.472) | 1.601(1.351-1.897) | 1.524(1.291-1.800) |
| 25-34 | 1.331(1.230-1.440) | 1.199(1.069-1.344) | 1.189(1.041-1.358) |
| 35-49 | Reference |  |  |
| **Marital Status** |  |  |  |
| Currently Married | Reference |  |  |
| Widowed | 1.078(0.902-1.289) | 1.177(0.961-1.441) | 0.671(0.484-0.930) |
| Separated/Divorced | 3.467(3.022-3.979) | 1.847(1.584-2.154) | 1.747(1.469-2.077) |
| **Residence** |  |  |  |
| Urban | 1.138(1.059-1.223) | 0.879(0.787-0.981) | 1.203(1.068-1.354) |
| **Education** |  |  |  |
| No education | Reference |  |  |
| Primary | 2.568(2.344-2.815) | 1.325(1.148-1.531) | 1.183(1.043-1.342) |
| Secondary | 2.461(2.250-2.692) | 1.049(0.880-1.250) | 1.258(1.039-1.523) |
| Tertiary | 1.282(1.090-1.507) | 0.667(0.492-0.903) | 0.839(0.461-1.528) |
| **Husband education** |  |  |  |
| No education | Reference |  |  |
| Primary | 2.539(2.292-2.813) | 1.367(1.170-1.598) | 1.157(0.997-1.342) |
| Secondary | 2.654(2.413-2.919) | 1.054(0.888-1.253) | 1.196(0.997-1.436) |
| Tertiary | 1.709(1.509-1.936) | 0.719(0.559-0.924) | 0.691(0.416-1.145) |
| **Employment** |  |  |  |
| Employed | 1.452(1.327-1.590) | 1.453(1.283-1.646) | 1.249(1.113-1.403) |
| **Husband employment** |  |  |  |
| Employed | 0.964(0.709-1.312) | 1.202(0.809-1.785) | 1.162(0.826-1.634) |
| **Wealth Category** |  |  |  |
| Poorest | Reference |  |  |
| Poorer | 1.399(1.240-1.577) | 1.256(1.090-1.448) | 0.813(0.673-0.983) |
| Middle | 1.772(1.577-1.991) | 1.101(0.947-1.280) | 0.811(0.673-0.977) |
| Richest | 1.673(1.487-1.882) | 0.900(0.769-1.053) | 0.949(0.792-1.137) |
| Richest | 1.587(1.406-1.791) | 0.647(0.536-0.782) | 0.998(0.836-1.192) |
| **Age difference** |  |  |  |
| Wife Older | Reference |  |  |
| Same age | 0.475(0.311-0.726) | 1.428(0.859-2.372) | 1.366(0.844-2.211) |
| Wife younger | 0.509(0.391-0.662) | 1.475(0.992-2.193) | 1.352(0.996-1.837) |
| **Husband takes alcohol** |  |  |  |
| Yes | 3.498(3.260-3.753) | 2.092(1.888-2.319) | 2.362(2.105-2.651) |
| **Husband Jealous** |  |  |  |
| Yes | 2.299(2.123-2.490) | 2.682(2.394-3.005) | 2.284(2.017-2.588) |

HR Hazard Ratio CI Confidence Interval

In Nigeria, ever-married women living in urban areas have 14% higher risk of experiencing domestic violence compared to their rural counterparts (HR=1.138: 95% CI=1.059-1.232). Compared with ever-married women with no formal education, the risk of domestic violence increased with increasing level of educational attainment except among those that had higher education in the three countries. The risks were 2.6, 2.5 and 1.3 times higher among ever-married women with primary, secondary and higher education in Nigeria respectively (HR= 2.568; 95% CI=2.344-2.815; HR=2.461: 95% CI=2.250-2.692 and HR=1.282; 95% CI=1.090-1.507) respectively. The employed women in the three countries had significantly higher risks of domestic violence. For instance the risks were 1.5 times higher in Kenya and Nigeria and 1.2 times higher in Mozambique. The risk of first domestic violence appeared to increase with higher wealth quintiles in Nigeria but the reverse was the case in Mozambique while women from households in the richest wealth quintiles in Kenya had 35% times lower risk of experiencing domestic violence compared with those in the poorest category (HR=0.647: 95% CI=0.536-0.782). Women whose husbands take alcohol and get jealous when the woman talks to another man were more than double likely to first have the risk of domestic violence than others across the three countries.

Table 4 shows the outcome of the multiple Cox proportional hazard model of the timing of first domestic violence in the three countries while controlling for confounders. The adjusted risk of first domestic violence after marriage was about 1.7, 1.5, and 1.6 times higher among ever-married women aged 15-19 years than women aged 45-49 years in Nigeria, Kenya and Mozambique respectively. Ever-married women aged 20-24 years in Kenya were about twice likely to fall a victim of domestic violence earlier (HR=1.937; 95% CI=1.586-2.366) compared to age group 35-49 years. The likelihood of an ever-married woman to have an earlier experience of domestic violence was significantly higher among separated/divorced women across the three countries than other women. (H.R=2.424: 95% CI=1.962-2.995; HR=1.266: 95% CI=1.006-1.594, and HR=1.445: 95% CI=1.106-1.887 for Nigeria, Kenya and Mozambique respectively).

**Table 4:** The Adjusted determinants of Violence after marriage.

| Characteristics | Adjusted determinants of Violence |  |  |
| --- | --- | --- | --- |
|  | Nigeria | Kenya | Mozambique |
| **Age** |  |  |  |
| 15-19 | 1.742(1.374-2.209) | 1.544(0.862-2.766) | 1.621(1.093-2.405) |
| 20-24 | 1.454(1.269-1.665) | 1.937(1.586-2.366) | 1.493(1.146-1.945) |
| 25-34 | 1.366(1.249-1.494) | 1.248(1.087-1.432) | 1.181(0.954-1.461) |
| 35-49 |  |  |  |
| **Marital Status** |  |  |  |
| Currently Married | Reference |  |  |
| Widowed | 1.012(0.849-1.208) | 0.844(0.691-1.030) | 1.196(0.936-1.528) |
| Separated/Divorced | 2.424(1.962-2.995) | 1.266(1.006-1.594) | 1.445(1.106-1.887) |
| **Residence** |  |  |  |
| Urban | 1.040(0.945-1.144) | 0.990(0.853-1.149) | 1.361(1.066-1.738) |
| **Education** |  |  |  |
| No education |  |  |  |
| Primary | 1.659(1.462-1.883) | 0.938(0.746-1.180) | 1.216(0.979-1.512) |
| Secondary | 1.579(1.371-1.818) | 0.951(0.723-1.251) | 1.226(0.802-1.874) |
| Tertiary | 1.126(0.902-1.406) | 0.826(0.525-1.298) | 1.567(0.553-4.442) |
| **Husband education** |  |  |  |
| No education | 1.650(1.440-1.891) | 1.102(0.868-1.399) | 1.047(0.821-1.336) |
| Primary | 1.674(1.453-1.929) | 0.829(0.631-1.090) | 0.944(0.661-1.349) |
| Secondary | 1.426(1.196-1.699) | 0.655(0.443-0.970) | 0.566(0.220-1.452) |
| Tertiary |  |  |  |
| **Employment** |  |  |  |
| Employed | 1.27(1.145-1.408) | 1.340(1.144-1.569) | 1.015(0.845-1.218) |
| **Husband Employed** |  |  |  |
| Employed | 1.034(0.72-1.485) | 1.141(0.682-1.908) | 0.657(0.424-1.017) |
| **Wealth Category** |  |  |  |
| Poorest |  |  |  |
| Poorer | 0.985(0.862-1.126) | 1.133(0.940-1.366) | 0.802(0.617-1.042) |
| Middle | 0.851(0.736-0.983) | 1.016(0.832-1.242) | 0.878(0.676-1.141) |
| Richest | 0.697(0.595-0.817) | 0.883(0.707-1.103) | 0.852(0.633-1.153) |
| Richest | 0.686(0.572-0.822) | 0.705(0.533-0.933) | 0.815(0.536-1.237) |
| **Age difference** |  |  |  |
| Wife Older | 0.495(0.316-0.777) | 1.806(0.998-3.269) | 1.010(0.475-2.150) |
| Same age | 0.581(0.440-0.765) | 1.680(1.039-2.718) | 1.201(0.745-1.934) |
| Wife younger |  |  |  |
| **Husband takes** |  |  |  |
| Yes | 2.789(2.557-3.042) | 1.936(1.708-2.194) | 2.615(2.176-3.144) |
| **Husband Jealous** |  |  |  |
| Yes | 2.381(2.180-2.600) | 2.252(1.976-2.567) | 2.493(2.038-3.050) |

aHR adjusted Hazard Ratio CI Confidence Interval Marital status dropped due to multicollinearity

The ever-married women with primary education had the highest risk of domestic violence in Nigeria, (HR=1.674: 95% CI=1.453-1.929) compared with those with no formal education but education was not a significant prognostic factor in Kenya and Mozambique. Furthermore, Nigerian women whose husbands had some education were generally more likely to experience domestic violence than those with no formal education, women with higher education had lower risks (HR=0.655: 95% CI=0.443-0.970) in Kenya but this was not significant in Mozambique. However, living in urban areas put women at higher risk of domestic violence.

Compared to the risks among ever-married women from households in the poorest wealth quintile, the richer the household from which a woman comes from the low in all the countries but this relationship was not significant in Mozambique. For instance women from richest households in Nigeria and Kenya had about 32% and 30% lesser risk of domestic violence than those from households in the poorest wealth quintiles respectively (HR=0.686: 95% CI=0.572-0.822 and HR=0.705; 95% CI=0.533-0.933). While employment status of the husbands were insignificant in the three countries, employed women in Nigeria and Kenya had higher risks of domestic violence than the unemployed women. Also the adjusted risks of first domestic violence was higher among women whose husbands take alcohol and get jealous when the woman talks to another man across board.

## DISCUSSION

While violence is a public health problem both in the developed and the developing countries. Females, particularly the younger girls, are the most vulnerable population subgroups since they are daily exposed and victimized by family members, at school, at work etc. It is more worrisome in the developing countries due to either non-availability of strict punishment or failure to implement the laws. Forced sex affects women of all ages across different socio-cultural boundaries. This study sought to determine the time at which women in Kenya, Zimbabwe and Cote d’Ivoire experience forced sex for the first time and the various factors influencing the timing of the first occurrence.

There is no sign that domestic violence is reducing in Africa. In fact, it seems to have defiled every intervention targeted at curtailing the menace in Africa. The prevalent socio-cultural belief in Africa that a man is the head of his family and is at liberty to discipline erring wife might have jeopardized efforts aimed at stalling domestic violence in sub-Sahara Africa. Domestic violence takes many forms including physical, sexual, emotional, and mental. While studies abate on the prevalence of domestic violence in marriages, there has been paucity of information on how soon a married women experience domestic violence after marriage. Neither has there been adequate information on the factors affecting the timing. The current study assessed the timings, identified the risk factors, compared the timings among those currently married and formerly married and also determined whether there are generational changes in the timing of first occurrence of domestic violence among ever-married women in three sub-Saharan countries.

We found that the timings of first violence were significantly influenced by the age of women (used as a proxy for birth cohorts), place of residence, women’s and husbands’ educational attainment and the households from which a woman comes from. Most dazzling among our finding is that more women born in the 1990s are experiencing domestic violence earlier in their marriage than those born 30 or 40 years earlier. We found a clear generational change in the time of first domestic violence among the cohorts of births with higher risk among younger ever-married women.

It is also a source of concern that the risk of domestic violence was significantly higher among the separated and divorced women than among the widows and among the currently married women. It has been reported in multiple studies that occurrence of violence against women is still high in the sub-Saharan Africa region, particularly in Nigeria which is the most populous country in Africa(12,13,22). Results from a recent study revealed that the total prevalence of spousal violence (physical, sexual or emotional) is about 30.5% in Nigeria 45.3% in Kenya and 45.5% in Mozambique(4).

The alarming rate of domestic violence in Africa remains an issue of concern. Despite the efforts geared toward reducing the prevalence, significant change has not been recorded over the years. In this study, the prevalence of domestic violence in Nigeria, Kenya and Mozambique are 12.3, 7.5%, and 15.9% respectively. This shows an insignificant improvement in unhealthy sociocultural norms that presupposes Nigerian women as inferior to men despite the development of gender policy in Nigeria to promote the health and rights of women.

The known associated factors responsible for high rate of domestic violence are age, education, employment, wealth status among others(19,31). The result of this study revealed that domestic violence was very evident among ever-married women of ages 15-19, 20-24, 25-29 and 30-34 years in Nigeria, Kenya and Mozambique. This is in consonance with a recent who multi-regional study which showed high prevalence of IPV among ever-married women across the earlier mentioned age groups. More importantly, this study agrees with the fact that IPV starts early in life (significantly high in age group 15-19) as suggested by the WHO multi-regional study(10). However, in this study, the highest report of domestic violence experience was found among respondents aged 20-24 (12%, 7% and 18% in Nigeria, Kenya and Mozambique respectively) as opposed to age 40-44(37.8%) in the multi-regional study. Similarly, a prior research showed that women younger than age 31are more probable to report past year spousal violence(31).

In our study, the association between violence and employment across the three countries was not striking enough, although highest violence prevalence was reported among women in the Agricultural sector while professional employment appeared to be protective of violence particularly in Mozambique. Prior studies have suggested that employment is significantly associated with the risk of domestic violence(32).

Besides, low or no literacy of women have been found by prior studies to be a predictor of violence while higher education, at least secondary school education have been found to be protective of violence(19,31,33,34). This is similar to what was found in this study - lower education of respondents and their partners seemed to increase the risk of violence across the three countries, suggesting that investing in education, especially women’s education may reduce the incidence of violence in sub-Saharan Africa. Notably, lower risks of violence was associated with higher education in Kenya, Mozambique and Nigeria. More so, higher education was found to be protective of violence in Kenya and Mozambique. However, respondents with secondary education had higher risk of violence than those with primary education in Nigeria. This mild difference may be indicative of cultural differences.

Another interesting finding is that rural residence was protective of violence among ever-married women in Nigeria. Whereas, in a study carried out in South India, higher prevalence of violence was associated with rural residence suggesting that the effect of location(urban and rural) on violence may be subject to regional differences(35). Furthermore, Women in the poorest wealth quintile had higher risk of violence in Nigeria, Kenya and Mozambique as opposed to prior multicountry study from sub-Saharan Africa which indicated that higher prevalence of violence was linked with higher household wealth quintile in Mozambique(4).

The observed level of domestic violence in this analysis might indicate that response to several campaigns against violence against women is still poor in African countries. Although, governments of different countries of Africa as the case in Nigeria have made several policies to curb violence against women, it seems the implementation of such policies is still far below the expectation. Even the calls by several non-governmental organizations advocating for “no violence against women” do not appear being effective yet. It is therefore important for policy makers to design a well implemented interventions that will promote peaceful cohabitations among domestics in Africa.

Our findings have some limitations. The census cannot be entirely replaced by the NDHS data and thus, there might still be some variations in the observed response. The question about the first experience of violence after marriage is prone to recall bias and sometimes, the information provided may not be absolutely correct. It would also have been useful to analyze the role of respondents’ behavior in the action of their partners against them. Although never-married women were excluded in the analysis, this exclusion is nevertheless not suggestive of nonexistence of physical or sexual violence among never-married women.

Despite all these limitations, this study has a significant message for policy against domestic violence in Africa. Firstly, the analysis clearly showed that a large number of women in Nigeria, Kenya and Mozambique are experiencing violence after marriage, which has led to dissolution of unions, disabilities and even deaths in many instances. Secondly, the residential variations in the experience of violence and other relevant factors will assist policy makers in identifying gaps in current programs geared towards preaching against violence in marriages and also guide development partners to identify critical groups among the ever-married women for their interventions.

The disparities in the effect of household income on the prevalence of domestic violence in Mozambique and Kenya is an indication that addressing household-poverty may not automatically aid the elimination of domestic in sub-Saharan(4), rather a wholesome approach must be taken to combat the menace.

## CONCLUSION

Domestic violence was quite high among ever-married women in Nigeria, Kenya and Mozambique. Higher prevalence of violence was among those no longer living with partner compared to those still with partner. Different socio-demographic characteristics (of the victim and the perpetrator) are directly associated with the problem, especially, education, age and wealth quintile. We therefore recommend that policy makers, government and other relevant stakeholders from the private sector should establish effective strategies towards minimizing the identified risk factors in order to reduce and prevent spousal violence as well as promote the health and wellbeing of women.

### Ethical approval

Ethical approval was sought from the National Health Research ethics committees in all the three countries by the data originators and granted before starting the survey. Also, informed consents were received from the participants before interviewing them. We obtained the approval from Measure of DHS for permission to use the data before analysis.

